# A sequence-based method for predicting extant fold switchers that undergo α-helix <-> β-strand transitions

**DOI:** 10.1101/2021.01.14.426714

**Authors:** Soumya Mishra, Loren L. Looger, Lauren L. Porter

## Abstract

Extant fold-switching proteins remodel their secondary structures and change their functions in response to cellular stimuli, regulating biological processes and affecting human health. In spite of their biological importance, these proteins remain understudied. Few representative examples of fold switchers are available in the Protein Data Bank, and they are difficult to predict. In fact, all 96 experimentally validated examples of extant fold switchers were stumbled upon by chance. Thus, predictive methods are needed to expedite the process of discovering and characterizing more of these shapeshifting proteins. Previous approaches require a solved structure or all-atom simulations, greatly constraining their use. Here, we propose a high-throughput sequence-based method for predicting extant fold switchers that transition from α-helix in one conformation to β-strand in the other. This method leverages two previous observations: (1) α-helix <-> β-strand prediction discrepancies from JPred4 are a robust predictor of fold switching, and (2) the fold-switching regions (FSRs) of some extant fold switchers have different secondary structure propensities when expressed in isolation (isolated FSRs) than when expressed within the context of their parent protein (contextualized FSRs). Combining these two observations, we ran JPred4 on the sequences of isolated and contextualized FSRs from 14 known extant fold switchers and found α-helix <->β-strand prediction discrepancies in every case. To test the overall robustness of this finding, we randomly selected regions of proteins not expected to switch folds (single-fold proteins) and found significantly fewer α-helix <-> β-strand prediction discrepancies (p < 4.2*10^−20^, Kolmogorov-Smirnov test). Combining these discrepancies with the overall percentage of predicted secondary structure, we developed a classifier that often robustly identifies extant fold switchers (Matthews Correlation Coefficient of 0.70). Although this classifier had a high false negative rate (6/14), its false positive rate was very low (1/211), suggesting that it can be used to predict a subset of extant fold switchers from billions of available genomic sequences.

## 1. Introduction

Extant fold-switching proteins remodel their secondary structures and change their functions in response to cellular stimuli^1^. Known instances of these environmentally responsive shapeshifters perform over 30 diverse functions, occur in all domains of life, and are associated with diseases such as cancer^2^, autoimmune disorders^3^, and malaria^4^. Furthermore, increasing evidence suggests that extant fold switchers are central to the regulation of biological processes^5^ such as cyanobacterial circadian rhythms^6^ and transcription/translation of bacterial virulence genes^7^.

Compared with single-fold proteins, which maintain stable secondary and tertiary structures and typically perform one biological function, extant fold switchers are understudied. Specifically, out of the ∼160,000 proteins with solved structures available in the Protein Data Bank (PDB^8^), fewer than 100 have been shown to switch folds. Increasing evidence suggests that fold switching is likely more widespread than currently appreciated^1^, but the current shortage of experimental examples makes it difficult to determine either the physical chemical properties or the functional scope of fold switchers. Thus, predictive tools are needed to identify more.

Recent computational studies suggest that fold switching is predictable, a prospect that—if realized—could greatly expand the small pool of experimentally determined fold switchers currently available. For example, naturally occurring extant fold switchers were predicted blindly by searching for differences between predicted and experimentally determined protein structures ^1,9^. Furthermore, a couple of fold-switching proteins have been designed computationally using the Rosetta software suite^10, 11^. Finally, progress has been made in predicting mutation-induced fold switching^12, 13^ as well as other conformational changes, such as rigid body motions^14^.

To date, all computational methods used to predict and design extant fold switchers suffer from the same drawback: the need for a protein structure on which to base predictions. This drawback severely limits the number of proteins that can be assessed for fold switching. Specifically, there are nearly 190,000,000 non-redundant, well-annotated protein sequences available in the RefSeq database^15^ but only 160,000 protein structures available in the PDB– which group into <15,000 unique structures^16^. Thus, any method that requires a solved protein structure to make predictions can sample < 0.01-0.1% of the protein sequences available. Furthermore, methods like Rosetta perform atomistic simulations, which are computationally intensive and not suitable for predicting the structures of thousands of proteins in parallel. For these reasons, sequence-based methods are needed to effectively identify fold switchers from the wealth of available genomic sequences.

Here we present a sequence-based method for predicting extant fold switchers that is based on the following hypothesis: the JPred4 secondary structure prediction of an isolated fold-switching region (FSR) sequence might differ from the JPred4 prediction of the same FSR within the context of its naturally occurring sequence (hereafter called a contextualized FSR). The basis for this hypothesis follows. We started from the previous observation^1^ that extant fold-switching proteins generally have: (1) regions that change secondary structure between the two forms (FSRs) and (2) regions that maintain the same secondary structure (structurally constant regions, or SCRs^17^). By definition, FSRs assume multiple stable secondary structures, and several studies have suggested that at least one FSR conformation is stabilized by exogenous interactions^18, 19^. Together, these observations indicate that the dominant secondary structure of a given FSR might differ depending on the context of its sequence. Thus, an FSR might assume a different secondary structure when expressed in isolation than when expressed within a larger protein sequence. Indeed, this is known to be the case for the C-terminal domain of RfaH, which folds into an α-helical bundle when covalently attached to its N-terminal domain but assumes a β-barrel conformation when expressed as an isolated domain^20^.

Since previous work strongly suggests that fold switching is most robustly predicted using α-helix <-> β-sheet prediction discrepancies from JPred4^13, 21^, we tested our hypothesis on 14 extant fold switchers with substantial α-helix <-> β-sheet transitions. In all 14 cases, JPred4 predicted that isolated and contextualized FSR sequences would have different α-helix and β-strand content. Specifically, overlapping regions were predicted to be α-helix for the isolated/contextualized sequence and β-sheet for the other sequence. To test the robustness of this finding, we randomly selected regions within a set of 211 proteins not expected to switch folds and found the secondary structure predictions of isolated and contextualized sequences to be significantly more consistent than they were for fold-switching proteins (p < 4.2*10^−20^, Kolmogorov-Smirnov test). We used this result to develop a classifier for extant fold switchers, which proved to be robust (Matthews Correlation Coefficient = 0.70). These results suggest that our approach can be used to predict a subset of extant fold switchers from the broad base of available genomic sequences.

## 2. Methods

### 2.1. Selection of extant fold switchers

Using a previous dataset^1^, we selected all extant fold switchers with FSRs ≥ 20 residues (the minimum length for JPred4 predictions) and an FSR region with at least 4 contiguous residues that assume an α-helix in one conformation and a β-strand in the other. This corresponded to a minimum secondary structure difference score of 0.6. (Secondary structure differences were scored as previously: α-helix <-> β-strand: 1; coil <-> α-helix or coil <-> β-strand: 0.5; no change in secondary structure: 0. Overall score is the total score normalized by FSR length.) Secondary structures of all extant fold switchers were classified using DSSP^22^.

### 2.2. JPred4 predictions of extant fold switchers

All amino acid sequences from 14 extant fold switchers with solved structures were downloaded from the Protein Data Bank (PDB) and saved as individual FASTA files. JPred4 predictions were run remotely using a publicly downloadable scheduler available on the JPred4 website (http://compbio.dundee.ac.uk/jpred/), and jnetpred predictions were used for all calculations. Each residue was assigned one of three secondary structures: “H” for helix, “E” for extended β-strand, and “C” for coil. Chain breaks were annotated “-”. PDB IDs and chains of each fold-switched pair were as follows: PimA_open_/PimA_closed_: 4NC9C/4N9WA; Selecase_4_/Selecase_1_: 4QHHA/4QHFA; RfaH_transcription_/RfaH_translation_: 2OUGA, 2LCLA; KaiB-_1_/KaiB_4_ (4 variants): 5JYTA/2QKEA,1WWJA,4KSOA,1R5PA; PfPRS_apo_/PfPRS_liganded_: 4YDQA/4TWAA; Ovalbumin_cleaved_/Ovalbumin_uncleaved_: 1JTIB/1OVAA; Proplasmepsin/Plasmepsin: 1MIQA/1QS8B; SUN2/SUN2-KASH1: 4DXTA/4DXRA; Aβ fibril/Aβ peptide: 2NAOF/1ITYA; Amylin/Amylin fibril: 2KB8A/6VW2A; α-synuclein fibril/α-synuclein: 2N0AD/2KKWA. All naturally occurring KaiB variants from cyanobacteria were included because we presumed that they all switch folds, though this has only been shown explicitly for *S. elongatus*. One additional hit (human primase—3L9Q/4RR2) was excluded because very recent NMR work strongly suggests that it is not a fold switcher but rather maintains its helical structure in solution; the alternative β-hairpin conformation is likely a crystallographic artifact^23^. FSR boundaries were initially chosen based on the regions reported previously (bold sequences in Table S2 of^1^). Fragments of KaiB and PimA were shortened by 13 and 17 residues, respectively, to yield secondary structure prediction discrepancies (sequences in Table S1). An additional 11 residues were also added to the N-terminal end of PimA’s FSR. Such modifications seemed reasonable since JPred4 makes predictions based on a 20-residue window^21^ that it could use to associate an isolated fragment with its contextualized secondary structure prediction. Thus, modifying short stretches of N- and C-terminal sequence could decrease the association between isolated sequences and their contextualized predictions.

### 2.3. Single-fold proteins and fragments

Proteins expected not to switch folds and having fewer than 800 residues (the upper limit in JPred4), totaling 211, were taken from Table S3C of^1^. One segment was selected from a random region of each protein. Segment lengths were randomly selected from the distribution of 14 FSR lengths (the lengths of the sequences in **Table S1**). Random selections were performed using the *random* module of Python2.7. JPred4 was run on all 422 sequences (211 full sequences + 211 segments) using its mass submit scheduler (http://www.compbio.dundee.ac.uk/jpred4/api.shtml#massSubmit).

### 2.4 Helix <-> strand discrepancies and distribution

Sequences of isolated FSRs were aligned with full-length proteins using the pairwise2.align.localxs function from Biopython^24^ with gap open/extension penalties of −1.0/- 0.5. Secondary structure predictions were re-registered according to the resulting alignments and compared. Helix <-> strand discrepancies between the predictions were summed residue-by-residue (1 for discrepancy, 0 for no discrepancy) and normalized by the minimum number of secondary structure annotations (helix and sheet) in the pair of sequences compared.

### 2.5. Distributions and statistics

The distributions in **Figures 2 and 3** was generated with Matplotlib^25^. Matthews Correlation Coefficients^26^ were calculated as follows:

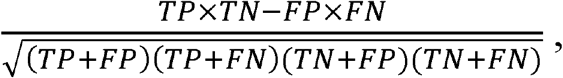

where TP = number of true positives, TN = number of true negatives, FP = number of false positives, and FN is the number of false negatives.

## 3. Results

3.1. Extant fold switchers with sizeable α-helix <->β-sheet transitions

We searched a previously published set of fold-switching proteins for all instances in which FSRs were at least 20 residues long and had at least four contiguous residues forming an α-helix in one conformation formed a β-strand in the other. This yielded 11 unique proteins (**Figure 1**), each which are highlighted briefly:

**Figure 1.**
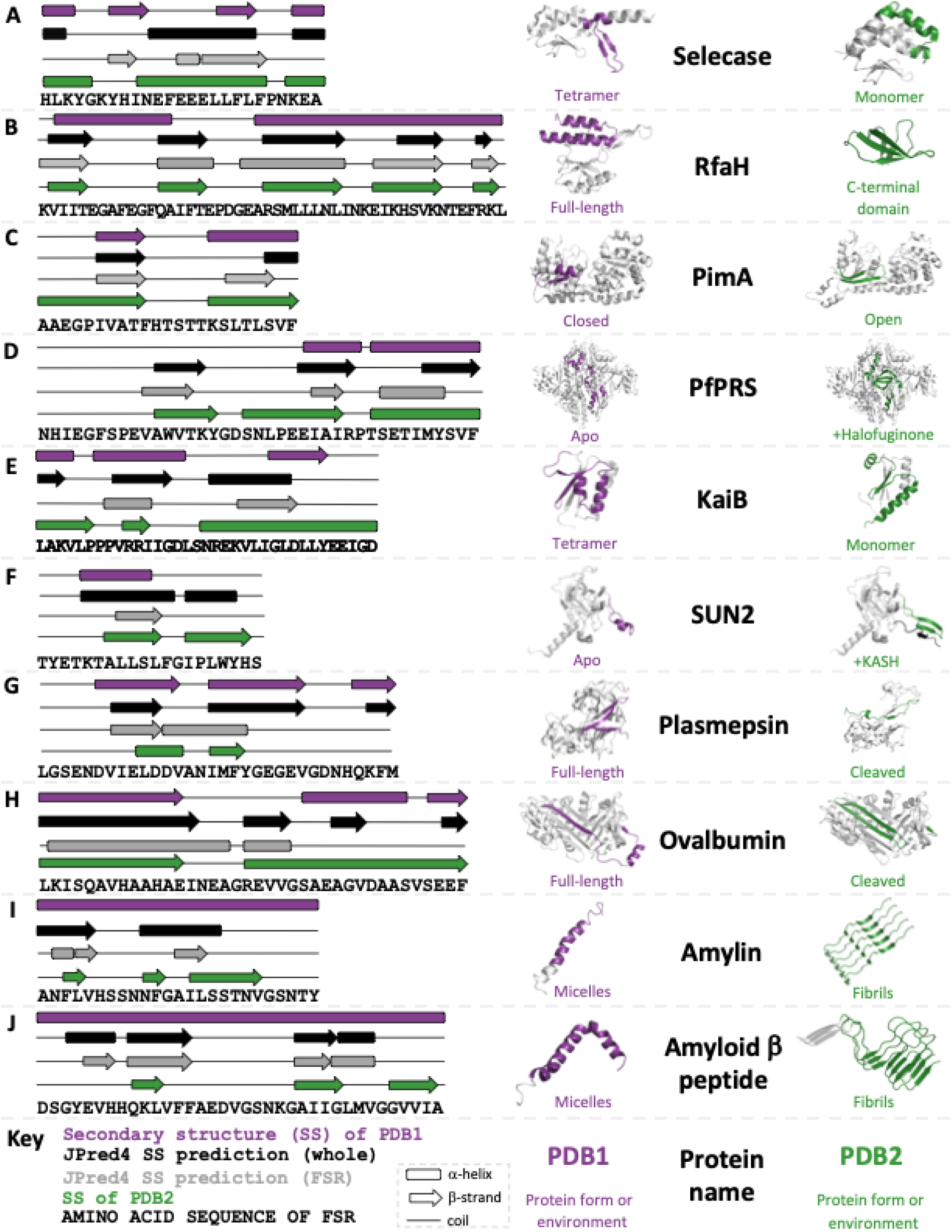
JPred4 predicts different secondary structures for isolated and contextualized FSRs of extant fold switchers with substantive transitions between α-helix and β-strand. Each panel shows the experimentally determined secondary structures of both conformations of the fold switcher (purple and green) along with JPred4 secondary structure predictions of the whole sequence (black) and FSR (gray). Purple and green regions of protein structure correspond to FSR sequence shown in diagram; gray do not. Predicted secondary structures that were at least 2 contiguous residues long are shown. The KaiB variant (2QKE) represents all members of the KaiB family; the other 3 (1WWJ, 4SKO, 1R5P) are not shown; α-Synuclein is also not shown due to lack of space. The C-terminus of KASH (SUN2 panel, green structure) is shown in black. All three-dimensional protein structures were made using PyMOL^48^.

- Selecase (“selective and specific caseinolytic metallopeptidase”; **Figure 1A**), produced by archaea and bacteria, and most studied from the archaeon *Methanocaldococcus jannaschii*, is an active metallopeptidase in its monomeric form. Upon forming structured higher order oligomers, namely dimers, tetramers, and octamers, Selecase is inactivated^27^. Its structures and activities are regulated by its concentration: mostly monomers at 0-0.3 mg/ml; dimers at 0.3-2 mg/ml; tetramers at 2-6 mg/ml, and octamers at > 6 mg/ml.
- RfaH (**Figure 1B**) regulates the expression of virulence proteins from enterobacteria such as *Escherichia coli*^*20*^. It has two domains: an N-terminal NGN-binding domain (NTD) and a C-terminal domain (CTD) that switches folds. RfaH’s CTD folds into an α-helical bundle that forms a binding interface with the NTD, masking its RNA polymerase (RNAP) binding site. Upon binding both RNAP and a specific DNA consensus sequence, called *ops*, the CTD dissociates from the NTD, unmasking the NTD’s RNAP binding site. This binding event also triggers the CTD to reversibly refold into a β-barrel able to bind the integral S10 unit of the ribosome and foster efficient translation^28^. When expressed in isolation, RfaH’s CTD folds into a β-barrel with no trace of α-helical content (green structure)^28^.
- PimA (**Figure 1C**) is a membrane-associated bacterial glycosyltransferase (phosphatidyl-*myo*-inositol mannosyltransferase) that initiates the biosynthesis of virulence factors produced by *Mycobacterium tuberculosis*. This enzyme has both a closed GDP-bound form and an open form with reshuffled secondary structure. PimA’s FSR is highly conserved in mycobacterial orthologs, and both crystallographic and near-UV CD evidence indicate that its open form could play an important role in membrane interactions^29^.
- PfPRS (**Figure 1D**) is a homodimeric prolyl tRNA synthetase from the malaria parasite *Plasmodium falciparum*. When bound to the inhibitor halofuginone, the active sites of the PfPRS monomers fold into β-sheets that hydrogen bond to active-site β-sheets of the other monomeric unit^30^. By contrast, the active site of apo PfPRS forms helical structures^4^.
- KaiB (**Figure 1E**) is a major component of the cyanobacterial circadian clock of *Synechococcus elongatus*^*6*^. Unlike most other circadian clocks, which are driven by transcription-translation oscillation, the cyanobacterial circadian clock is maintained through a periodic phosphorylation cycle, known as a post-translational oscillator (PTO)^31^. At night, KaiB’s active monomeric form helps to regulate the dephosphorylation of the PTO, while in the morning it primarily populates an inactive tetramer with a different fold, allowing phosphorylation of the PTO.
- SUN2 (**Figure 1F**), in complex with its binding partner KASH, tethers the nuclear envelope to the cytoskeleton, making it critical to meiosis, nuclear migration, and global cytoskeletal organization^32^. A region of SUN2, known as the KASH lid, is likely flexible in apo SUN2 and forms a small helix in the apo crystal structure. By contrast, the KASH lid forms a hydrogen-bonded β-sheet with KASH’s C-terminal segment in the complex (black). Both mutational and functional analyses highlight the importance of the SUN-KASH β-sheet: alanine mutations at conserved positions within the β-sheet of both SUN and KASH decreased binding significantly, and deletion of the KASH lid compromised targeting to the nuclear envelope^32^.
- The pathogenic malarial aspartic protease plasmepsin (a family of at least 10 closely related enzymes in *P. falciparum*^*33*^; **Figure 1G**) degrades human hemoglobin and other globins – producing many of the disease’s symptoms^34^. It has a zymogenic form with an N-terminal propeptide that targets it to the protist’s food vacuole, while also forcing the protein into a conformation that inhibits its activity. A maturase activates plasmepsin by cleaving its propeptide, the removal of which allows its N-terminus to dramatically refold into a conformation capable of proteolytic activity^34^.
- Ovalbumin (**Figure 1H**) is a member of the serpin family (serine protease inhibitor; although ovalbumin is not known to have *in situ* inhibitory activity – it constitutes 60-65% of egg whites and appears to be a storage protein^35^) with a zymogenic form (*i*.*e*., an inactive precursor, as has plasmepsin). Specifically, inactive ovalbumin has a reactive center loop (RCL) that, when cleaved by a serine protease such as subtilisin, forms a β-strand inserted between two pairs of β-hairpins on its surface. Additionally, the α-helix formed by ovalbumin’s uncleaved RCL is regular and less flexible than the distorted helices of inhibitory serpins^36^.
- The human amyloid-forming peptides amylin and amyloid β (**Figures I & J**, respectively), along with α-Synuclein (Table S1) are all believed to interact with membranes, where they form α-helices^37-39^. While the cognate functions of helical α-synuclein and amyloid β remain under investigation, amylin is an endocrine hormone (co-secreted with insulin) that regulates glycemic metabolism^38^. All three peptides can also form fibrillar deposits associated with diseases such as Parkinson’s (α-synuclein)^40^, type 2 diabetes (amylin)^41^, and Alzheimer’s (amyloid β)^42^.

### 3.2. JPred4 predicts different secondary structures for isolated and contextualized FSRs

The proteins shown in **Figure 1** have fold-switching regions (FSRs, shown in color) and structurally constant regions (SCRs, shown in gray). Previous work suggests that the structures assumed by FSRs are context-dependent^18, 19^: that is, exogenous FSR-SCR interactions might bias the FSR away from its intrinsic secondary structure propensities, forcing it to adopt an alternative, higher-energy secondary structure conformation^20^. Thus, we sought to determine whether JPred4 might predict that FSRs can fold into multiple stable secondary structures by comparing the secondary structure predictions of isolated FSR sequences (shown in **Figure 1**) with those of the FSR sequence within its natural sequence context (hereafter called contextualized FSRs).

In all cases shown above, along with all other KaiB variants tested (**Table S1**), we found that JPred4 predicted different secondary structures for isolated and contextualized FSR sequences (**Figure 1**). Furthermore, in 12/14 cases, the regions of α-helix <-> β-strand prediction discrepancy (where JPred4 predicted α-helix for an FSR sequence in one context and a β-strand for the same FSR in its other context) corresponded to a region where the sequence indeed assumes an α-helix in one experimentally determined conformation and a β-strand in the other. In the two cases where this did not occur (PfPRS and Ovalbumin, **Figure 1D and 1G**, respectively), one region of α-helix <-> β-strand prediction discrepancy was within 5 residues of an experimentally determined FSR that assumed an α-helix in one conformation and a β-strand in the other.

JPred4 secondary structure predictions tend to correspond reasonably well (<6% α-helix <-> β-strand discrepancies^9^) with at least one experimentally determined protein structure for all 14 proteins. In fact, in 10 of these cases, α-helix and β-strand secondary structure elements correspond well between one prediction and one experimentally determined conformation (correct secondary structures in all of the right positions, though not necessarily the experimentally determined lengths). However, JPred4 predictions correspond well to both experimentally determined conformations in only 2/14 cases. Nevertheless, as in previous work^13^, we use discrepancies between predictions to infer fold switching; for our purposes the accuracies of the JPred4 predictions have no bearing on this inference.

### 3.3. JPred4’s α-helix <-> β-strand prediction discrepancies are statistically significant for FSRs

We then sought to determine the significance of JPred4’s α-helix <-> β-strand prediction discrepancies for isolated and contextualized FSRs. To do this, we randomly selected fragments from a set of 211 proteins expected not to switch folds (single-fold proteins). The resulting distribution of α-helix <-> β-strand discrepancies for fragments of single-fold proteins differed significantly from that of extant fold switchers (p < 4.2*10^−20^, Kolmogorov-Smirnov test; **Figure 2**).

**Figure 2.**
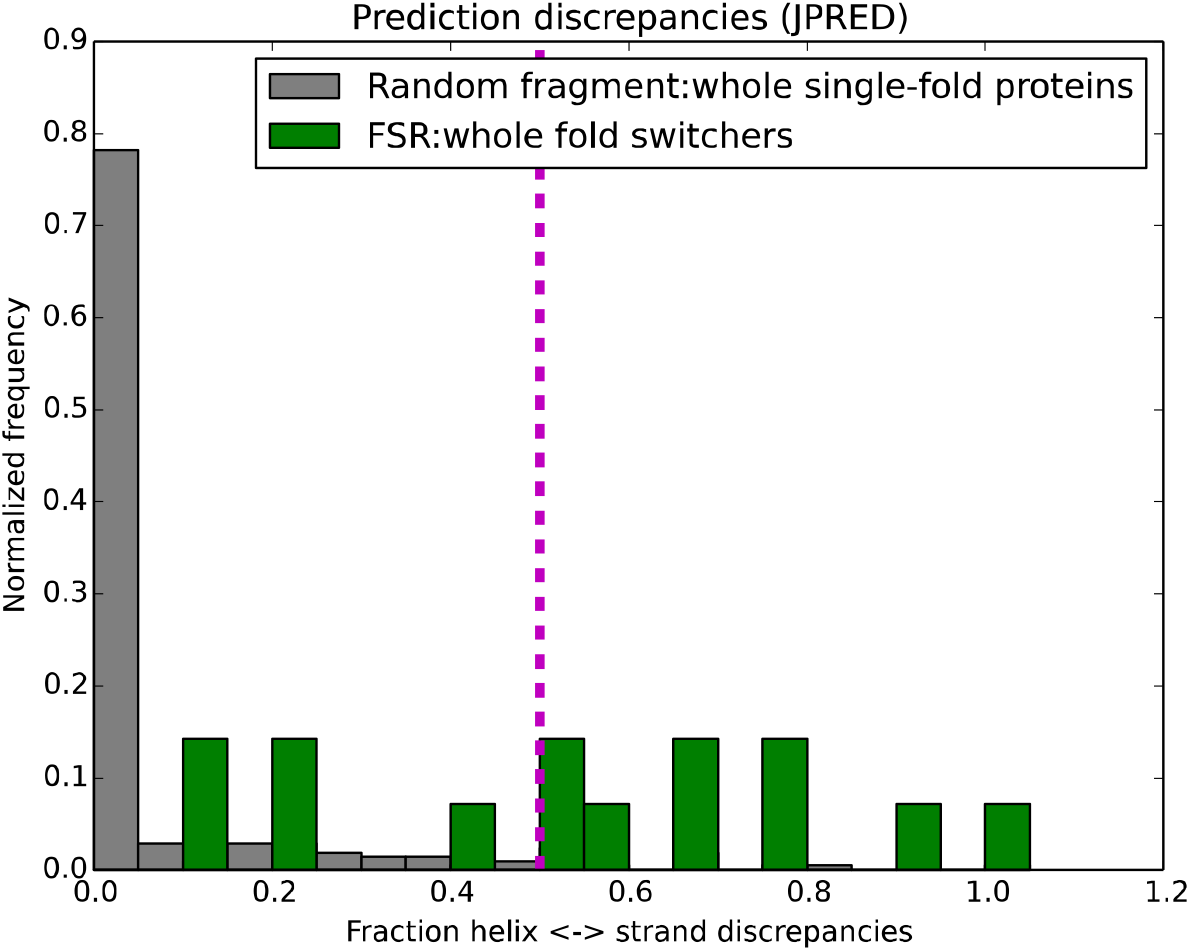
The distributions of JPred4 α-helix <-> β-strand prediction discrepancies between fragment:whole sequence for single-fold proteins and fold switchers differ significantly (p < 4.2*10^−20^, Kolmogorov-Smirnov test). Magenta dashed line represents the classifier threshold of 0.5, which yields a Matthews Correlation Coefficient of 0.47.

We then used the fraction of α-helix <->β-strand discrepancies as a potential classifier for predicting fold switchers. At a threshold of 0.5 (total number of helix <->strand discrepancies normalized by total number of residues predicted to assume regular secondary structure, **Methods**), the Matthews Correlation Coefficient of this classifier was 0.47, demonstrating moderate agreement with experimental findings. With this constraint in place, this predictive method yielded 9 true positives, 5 false negatives, 198 true negatives, and 13 false positives. Thus, we sought for additional constraints to improve predictions.

### 3.4. Constraining by fraction of secondary structure decreases false positives

Many false positives identified by the previous classifier shared a common feature: low overall levels of predicted secondary structure (and, conversely, high overall levels of predicted coil). To quantify the significance of this feature, we plotted the fraction of predicted secondary structure (total number of residues predicted to be helix or strand normalized by the total number of residues in the protein fragment) versus the fraction of secondary structure discrepancies for both FSRs and randomly selected regions of single-fold proteins (**Figure 3**). Indeed, we found that nearly all (12/13) false positives had low fractions of predicted secondary structure (<0.35).

**Figure 3.**
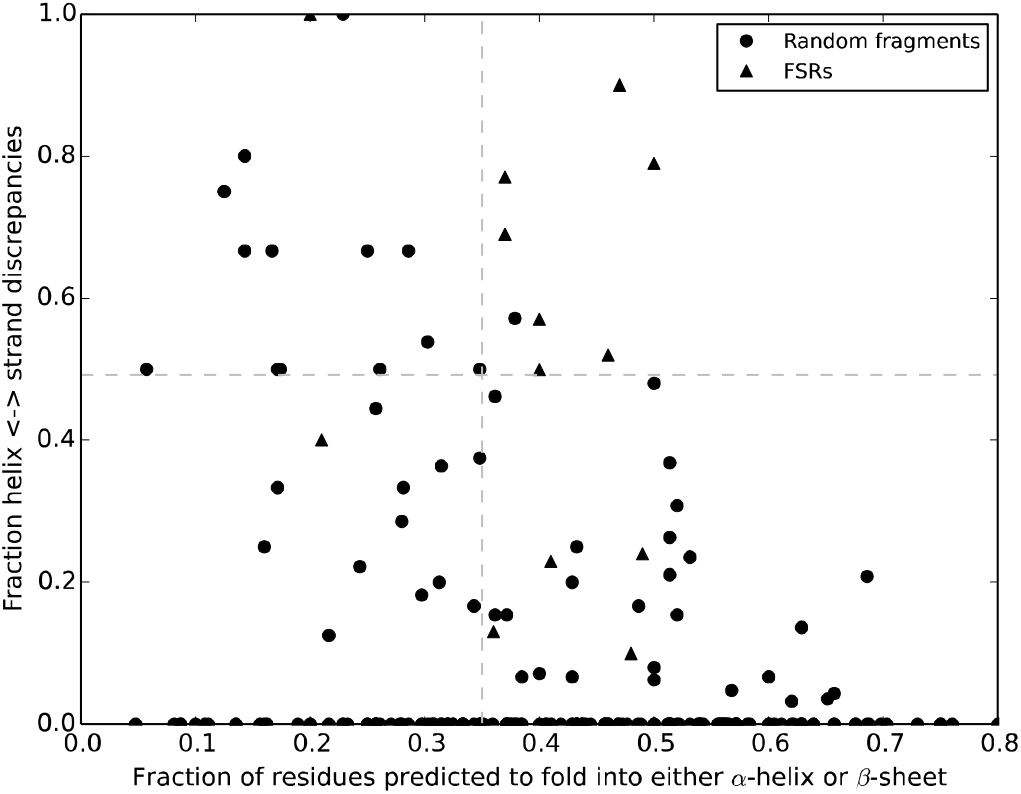
Predictions of extant fold switchers improve by constraining both the fraction of predicted secondary structure and the fraction of α-helix <-> β-strand discrepancies. Vertical/horizontal dashed lines represent the thresholds for fraction of predicted secondary structure/fraction of α-helix <-> β-strand discrepancies. Datapoints in the upper right quadrant are predicted to switch folds. Only 7/8 fold switchers can be seen because two KaiB variants have identical coordinates (0.37,0.69).

The one remaining false positive comprised residues 12-48 from the glutathione S-transferase (GST) Omega 3 expressed by the silkworm *Bombyx mori*. Residue 29 of this segment is an asparagine, which replaces a highly conserved cysteine in the other members of the Omega family^43^. This single amino acid change is partially responsible for Omega 3’s loss of GST activity: mutating asparagine 29 to a cysteine while also deleting its flexible C-terminal helix restores GST activity. Interestingly, running JPred4 on the same segment (residues 12-48) with just an N29C mutation gives the same secondary structure prediction as that of the whole protein (**Table S2**)^43^. Based on our previous work on sequence-similar fold switchers^13^, this result suggests that this protein segment might switch folds.

### 3.5. An improved classifier results from combining secondary structure prediction discrepancies with the fraction of secondary structure

Requiring a minimum of 35% secondary-structure content removed 12/13 false positives while eliminating only one true positive. Thus, we sought to classify fold switchers by thresholding our results by 0.5 for fraction of α-helix <->β-strand discrepancies and 0.35 by fraction of predicted secondary structure (**Figure 3**). This yielded 8 true positives, 1 false positive, 210 true negatives, and 6 false negatives, resulting in a Matthews Correlation Coefficient of 0.70 (very low false positive rate; moderate false negative rate).

## 4. Discussion

Fold switchers are exceptions to the well-supported general observation that folded proteins assume one stable structure that performs one function. Nevertheless, increasing evidence suggests that these proteins may be more abundant in nature than previously thought^1^. Fold switching impacts protein function^5^ and is associated with a number of diseases^2, 3, 30^. Thus, it would be useful to have a bioinformatic algorithm that identifies more fold switchers from their sequences. This is especially true because, up to this point, all experimentally characterized fold switchers have been stumbled upon by chance^1^.

Here we present an approach for predicting extant fold switchers from their amino acid sequences alone. To the best of our knowledge, this is the first sequence-based method for predicting extant fold switchers presented in the literature. This method is based on previous experimental work suggesting that the fold-switching regions (FSRs) of proteins are context-dependent: that is, their conformations are determined by their environment^18, 19^. In light of this, we hypothesized that it might be possible to predict extant fold switchers by comparing the JPred4 secondary structure predictions of isolated FSRs with contextualized FSRs and searching for α-helix <-> β-strand discrepancies. Indeed, such discrepancies were found in 14/14 fold switchers used in this study. Furthermore, this finding was highly statistically significant (p < 4.2*10^−20^, Kolmogorov-Smirnov test), and we used it to develop a classifier for extant fold switchers that yielded a Matthews Correlation Coefficient of 0.7. We suspect that JPred4 successfully identified extant fold switchers for the same reason it identified sequence-similar fold switchers^13^: different sequences (contextualized and isolated FSRs in this case) yielded different sequence profiles from PSI-BLAST searches. Future work revealing how these different profiles lead to dramatically different secondary structure predictions would be useful.

Two additional results stand out in light of previous work. First of all, the method presented here predicts fold switching in all four KaiB variants tested. This positive result is an improvement over our previous method for sequence-similar fold switchers, which failed to predict fold switching in all KaiB variants^13^. Secondly, our results strongly suggest that the fragment from Omega 3 is an FSR, even though it was in our set of proteins not expected to switch folds. Just one mutation (N29C) is sufficient to dramatically change the secondary structure predictions of this sequence, a previously identified characteristic of sequence-similar fold switchers (proteins with highly similar— but not identical—amino acid sequences and different folds^13^). Additionally, Omega 3’s GST topology^43^ has been known to switch folds in other proteins, namely KaiB^44^ and Chloride Intracellular Channel 1 (CLIC1)^45^. Still, further experimental work would be needed to determine whether or not Omega 3 switches folds.

Although we are optimistic that the approach presented here can be used to predict novel fold switchers, it has several limitations. Firstly, it can only identify fold switchers that undergo large α-helix <-> β-sheet transitions. To date, these proteins are rare and comprise only 15% of known fold switchers. Biologically important fold switchers similar to lymphotactin^46^, which maintains β-sheets that change their hydrogen bonding register, and β-pore proteins^47^, which extend already existing β-sheet structures, will be missed. Secondly, it will not identify all fold switchers that undergo large α-helix <-> β-sheet transitions, as evidenced by the fact that only 8/14 of the fold switchers tested gave a robust enough signal to be classified positively. Thirdly, because the FSRs of undiscovered fold switchers are not known *a priori*, our method will likely need to test many putative FSRs (different sizes and different regions) within the same protein to determine whether or not it is a fold switcher. Although this approach is much less computationally intensive than all-atom simulations, it will still require substantial time and computational power to predict fold switching in thousands of genomic sequences. Furthermore, the more sequences we probe, the more likely we are to hit false positives. Additional work will be needed to more accurately distinguish between these false positives and true fold switchers. Finally, our training set was small, comprising the only 14 fold switchers suitable for the predictive method presented here. Thus, it is likely that our statistics, especially for true positives and false negatives, are noisy. As more fold switchers are discovered, we are optimistic that it will be possible to develop methods that can predict more types of fold switchers with higher accuracy.

## 5. Conclusions

Our results suggest that the α-helix <-> β-strand transitions of some extant fold switchers can be predicted from their sequences alone using the homology-based secondary structure predictor JPred4. Although this method will not identify all extant fold switchers whose secondary structures transition from α-helix <-> β-strand, its low false positive (1/211) and moderate true positive (8/14) rates suggest that many positive predictions will likely correspond to true extant fold switchers. Thus, we are optimistic that this approach can be used to predict a subset of extant fold switchers from the broad base of available genomic sequences.

## Data availability

The data and code that support the findings of this study are openly available on GitHub at https://github.com/porterll/extant_fold_switchers

## Acknowledgements

This work utilized the computational resources of the NIH HPS Biowulf cluster (http://hpc.nih.gov). This work was supported in part by the Intramural Research Program of the National Library of Medicine, National Institutes of Health.

## Conflict of Interest Statement

The authors declare no conflict of interest.

---

**Table S1:**
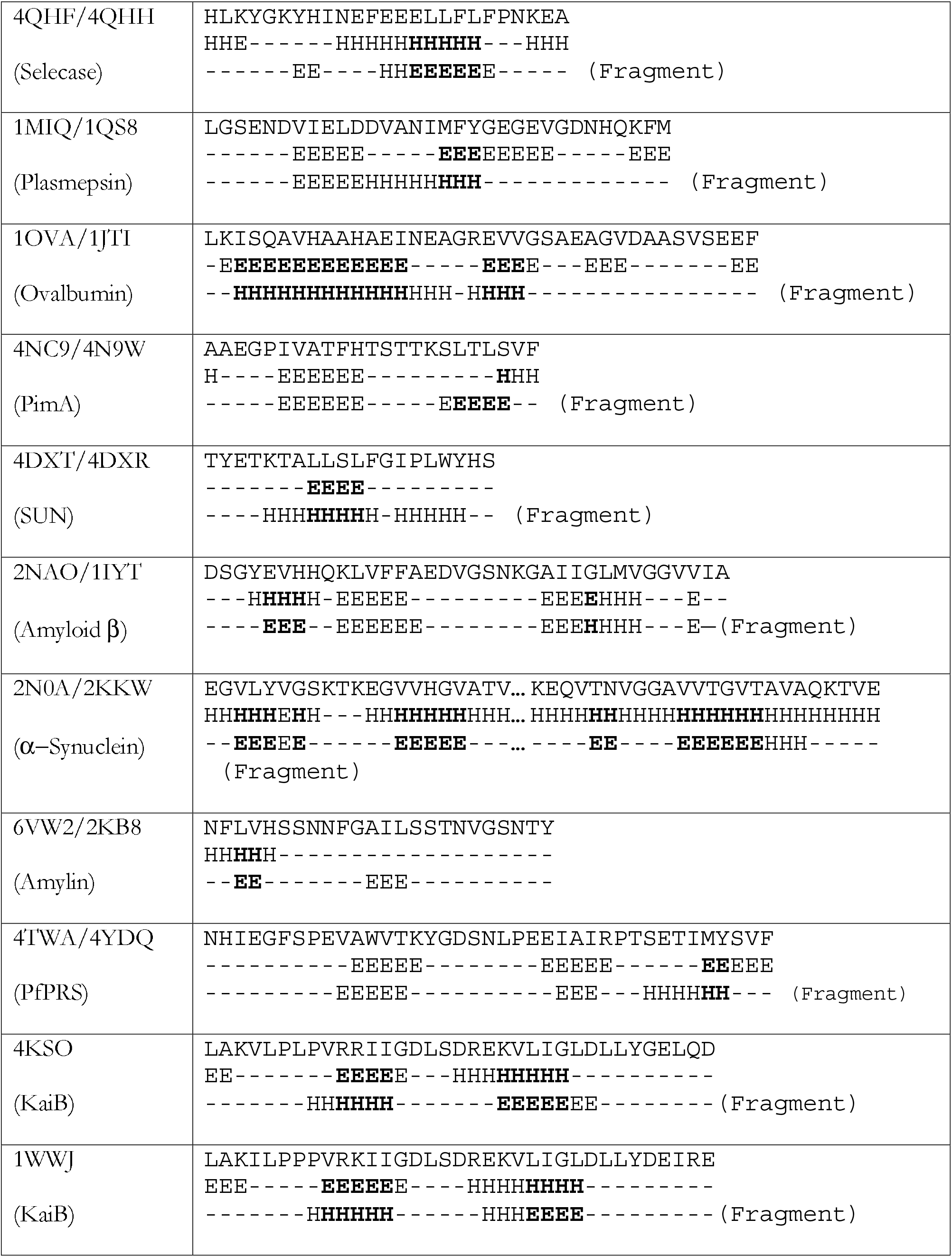
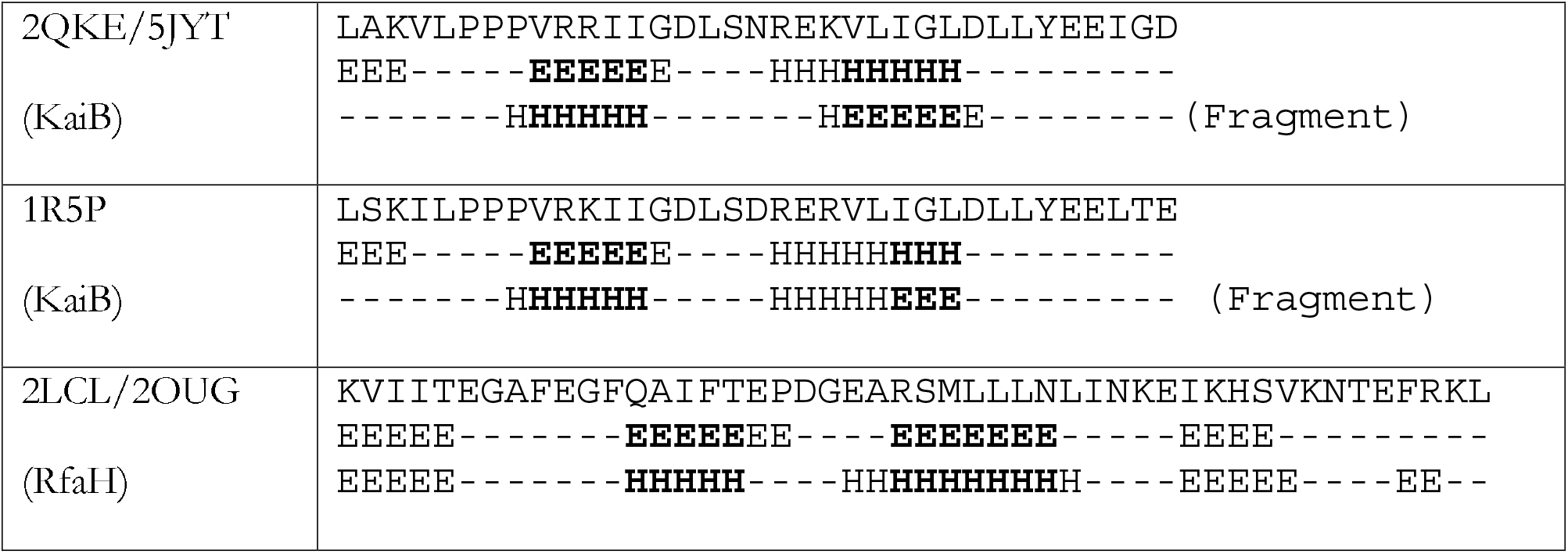
JPred4 predictions of extant fold switchers.

**Table S2.**
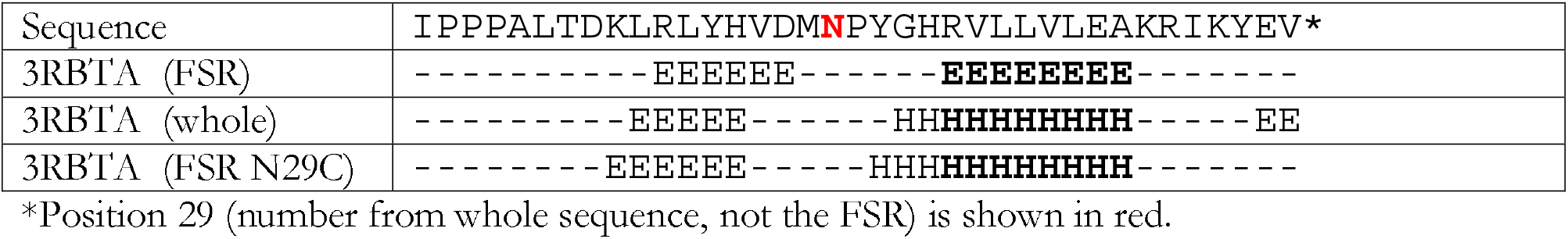

## Notes

### Competing Interest Statement

The authors have declared no competing interest.

